# An Inducible Model for Unraveling the Effects of Advanced Glycation End-Product Accumulation in Aging Connective Tissues

**DOI:** 10.1101/2020.09.04.283473

**Authors:** Austin G. Gouldin, M. Ethan Brown, Jennifer L. Puetzer

## Abstract

**Purpose:** In connective tissues there is a clear link between increasing age and degeneration. Advanced glycation end-products (AGEs) are believed to play a central role. AGEs are sugar-induced non-enzymatic crosslinks which accumulate in collagen with age and diabetes, altering tissue mechanics and cellular function. Despite ample correlative evidence linking collagen glycation to tissue degeneration, little is known how AGEs impact cell-matrix interactions, limiting therapeutic options. One reason for this limited understanding is AGEs are typically induced using high concentrations of ribose which decrease cell viability, making it impossible to investigate cell-matrix interactions. The objective of this study was to develop a system to trigger AGE accumulation while maintaining cell viability.

**Materials and Methods:** Using cell-seeded high density collagen gels, we investigated the effect of two systems for AGE induction, ribose at low concentrations (30, 100, and 200 mM) over 15 days of culture and riboflavin (0.25 mM and 0.75mM) induced with blue light for 40 seconds (riboflavin-465 nm).

**Results:** We found ribose and riboflavin-465 nm treatment produces fluorescent AGE quantities which match and/or exceed human fluorescent AGE levels for various tissues, ages, and diseases, without affecting cell viability or metabolism. Interestingly, a 40 second treatment of riboflavin-465 nm produced similar levels of fluorescent AGEs as 3 days of 100 mM ribose treatment.

**Conclusions:** Riboflavin-465 nm is a promising method to trigger AGE crosslinks on demand *in vivo* or *in vitro* without impacting cell viability and offers potential for unraveling the mechanism of AGEs in age and diabetes related tissue damage.

## Introduction

Generally, with aging and/or diabetes, tissues throughout the body, such as arteries,^1^ lungs,^2^ skin,^3^ lens,^4^ cartilage,^5^ tendons,^6^ and ligaments^6^ are known to become stiffer. This stiffening of the extracellular matrix leads to an increased rate of injury and degeneration of the tissues.^7^ In musculoskeletal tissues alone, 75% of individuals over the age of 65 will suffer an injury.^8^ While many mechanisms, such as aging stem cells, reactive oxidative species (ROS), and inflammation, contribute to this clear link between increasing age and degeneration, one of the most prominent mechanisms believed to be responsible, is progressive accumulation of advanced glycation end-products (AGEs).^7,9,10^ AGEs are sugar-induced non-enzymatic crosslinks that accumulate with age and/or diabetes in the extracellular matrix, altering tissue mechanics and cell function. AGEs were first identified in cooked food as end-products from a non-enzymatic reaction between sugars and proteins called the Maillard reaction.^11^ Around 40 years ago a similar glycation process was recognized in the extracellular matrix of diabetic patients.^12^ Since then it has been suggested that protein damage due to AGE crosslinking is the major cause of disease related pathogenesis in diabetes.^13,14^

The first step of AGE formation is initiated by covalent attachment of reducing sugars to amino groups of proteins, lipids, or nucleic acids to produce a reversible and unstable Schiff base.^11^ Then, the Schiff base may undergo Amadori rearrangement resulting in a more stable Amadori product. After a certain period of time, spanning from weeks to years, the Amadori product will transition to form an AGE crosslink or adduct.^7,9,15^ AGE formation is quite slow *in vivo*, therefore AGEs primarily form in long-lived proteins like collagen, and accumulate with age as cells decrease their turnover of the matrix.^16^ AGE crosslinks are thought to alter the mechanics of the ECM, as well as increase inflammation and oxidative stress through interaction with RAGE receptors, eventually leading to chronic illnesses.^12,14,17^ In fact, AGEs have been linked to many age-related diseases, such as atherosclerosis,^18^ chronic kidney disease,^19^ rheumatoid arthritis,^20–22^ and neurodegenerative diseases such as Alzheimer’s and Parkinson’s disease.^23^ Further, due to the high reliance on collagen in connective tissues such as tendon, ligament, meniscus, and cartilage, AGEs have been linked to age-related changes in mechanics that lead to increased injury and degeneration.^24–27^

AGEs are thought to induce disease and tissue damage by two main mechanisms. AGEs can either crosslink proteins, directly altering their structure and in-turn altering their properties and function, or alter binding sites on proteins changing cellular signaling.^12^ Despite ample correlative evidence connecting collagen glycation to aging and disease, little is known about how AGEs impact cell-matrix interactions, resulting in limited therapeutic options.^9,10^ It is difficult to study the AGE mechanism *in vivo* due to differences in collagen organization, disease state, and age.^9,10^ *In vitro* models are an attractive means to study AGEs because they eliminate these confounding variables. Traditionally AGE crosslinks have been induced through ribose culture in unorganized collagen gels and explanted rat tendons.^7,28,29^ Ribose is typically used to induce AGEs due to its higher reactivity rate, in comparison to glucose and fructose, which ultimately shortens the timeframe needed for AGE formation to occur.^30,31^ However, *in vitro* work is often performed with concentrations greater than 200 mM ribose, resulting in super-physiological levels of glycation and low cell viability due to hyperosmotic environments.^7,10,29,32^ Due to low cell viability in these systems it is impossible to study the cellular response to AGE protein modifications, which in turn greatly reduces the physiological reliability of the results. More specifically, a previous experiment exploring fibroblast viability in ribose treated collagen matrices concluded concentrations starting at 240 mM ribose lead to cell death.^29^ The objective of this study was to develop an inducible system to trigger native level AGEs *in vitro*, while maintaining cell viability, in order to study the effect of AGEs both on protein properties and cellular response.

One promising method for inducing AGEs instantaneously without cell viability issues is through riboflavin exposed to blue light (riboflavin-465 nm). Riboflavin, or B2 vitamin, is a non-toxic photosensitizer that when exposed to either UV light or blue light (465 nm) results in the formation of covalent crosslinks.^33^ Upon photoactivation, riboflavin will form covalent bonds with amino acids of collagen, as well as form reactive oxygen species that can further oxidize amino acids for reactions.^34^ The mechanism of riboflavin photoactivation is still not completely understood but the formation of covalent crosslinks between riboflavin and collagen is similar to the Maillard reaction in AGE formation.^33,35^ In addition, the process of generating oxygen free radicals while crosslinking collagen is a trend seen in AGE formation.^12,14,17^ Riboflavin is used clinically to enhance corneal strength using UV activation, and *in vitro* to increase the strength of different hydrogels.^33,36–42^ Specifically, it has been shown to crosslink collagen gels in a dose dependent fashion with limited viability issues, depending on the concentration of riboflavin and duration of light exposure.^38^ Riboflavin holds great potential to create an inducible AGE model; however, despite its wide use for crosslinking collagen gels, it has not been investigated as a means to induce AGEs. We hypothesize that riboflavin photoactivated by blue light in high density collagen gels will produce native fluorescent AGE levels without altering cell viability or metabolism. Additionally, in this study we evaluated traditional ribose methods of inducing AGEs, at lower concentrations typically reported in the literature, to investigate whether lower concentrations of ribose for longer periods of time can induce physiological levels of fluorescent AGE accumulation without altering cell viability or metabolism.

## Materials and Methods

### Cell-seeded construct fabrication

In this study, high density collagen gels (20 mg/ml) seeded with meniscal fibrochondrocytes were used as the construct system for AGE accumulation. Low-density collagen gels (1-5 mg/ml) have been traditionally used for collagen crosslink studies;^29,32,33,38,43,44^ however these gels often undergo significant contraction resulting in limited culture durations. High density collagen gels (10-20 mg/ml) have collagen concentrations closer to native values, reduced contraction compared to low density gels, and have been shown to maintain cell viability over 8 weeks of culture.^45–48^ Bovine meniscal fibrochondrocytes were used as a generic primary cell source, since menisci are substantially affected by AGE accumulation during aging^26,49^ and we have previously shown meniscal fibrochondrocytes survive in high density collagen gels for up to 8 weeks.^45,46,48,50^

#### Cell isolation

Bovine meniscal fibrochondrocytes were isolated from whole menisci within 24 hours of slaughter as previously reported.^45,46,48,50^ Briefly, 1-6 week old calf legs were purchased from a local slaughterhouse. Menisci were aseptically isolated from the joint, diced, and digested overnight in 0.3% collagenase (Worthington). The next day cells were filtered, washed, and counted. Cells from 3 donors were combined to limit donor variability. Cells were seeded at 2800 cells/cm^2^ into T-175 flasks and passaged 1-2 times to obtain enough cells. Cells were cultured in medium consisting of Dulbecco’s Modified Eagle Medium (DMEM), 10% fetal bovine serum (FBS), 1% Antibiotic antimycotic, 0.1 mM non-essential amino acids, 50 μg/mL ascorbic acid, and 0.8 mM L-proline.^45,46,48,50^

#### Construct fabrication

Cell seeded high density collagen gels were fabricated as previously described.^45–48^ Briefly type I collagen was extracted from purchased mix gender adult Sprague-Dawley rat tails (BIOIVT) and reconstituted at 30 mg/mL in 0.1% acetic acid.^45,46,48^ The stock collagen solution was mixed with appropriate volumes of 1 N NaOH, 10x phosphate buffer solution (PBS), and 1x PBS to begin the gelling process and return the collagen solution to pH 7 and 300 mOsm osmolarity.^47,51^ The collagen solution was immediately mixed with meniscal fibrochondrocytes suspended in media, injected between glass sheets 1.5 mm apart, and gelled at 37°C for 1 hour to obtain 20 mg/mL collagen sheet gels at 5×10^6^ cells/mL. Six mm biopsy punches were cut from sheet gels, split between experimental groups, and cultured for up to 15 days.

### Glycation culture conditions

After construct fabrication, cell-seeded constructs were split between DMEM control, riboflavin-465 nm, or ribose groups and cultured for up to 15 days (**Figure 1**). All constructs were maintained in standard incubator conditions at 37°C and 5% CO_2_, with medium renewal every 3 days. DMEM control constructs were cultured in standard growth media consisting of DMEM, 10% FBS, 1% Antibiotic antimycotic, 0.1 mM non-essential amino acids, 50 μg/mL ascorbic acid, and 0.8 mM L-proline. Constructs were collected for time points at day 0, 3, 6, 9, 12, and 15.

**Figure 1:**
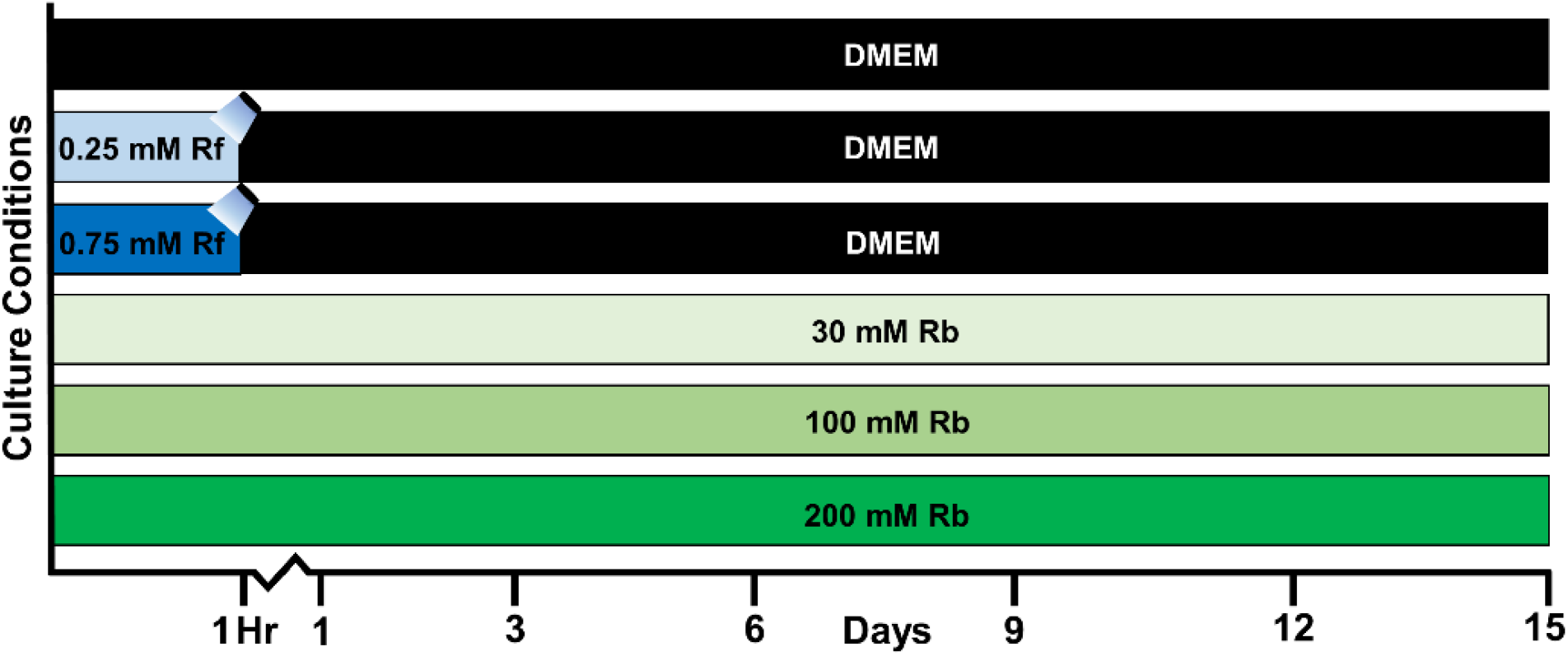
Exposure timeline for all control (DMEM), Riboflavin-465 nm (Rf), and Ribose (Rb) concentrations. Control constructs were kept in basic DMEM media for the full 15 days. Riboflavin-465 nm constructs were soaked in riboflavin for 1 hour, stimulated with blue light for 40 seconds to activate crosslinks and then cultured in basic DMEM for 15 days. Ribose constructs were cultured in basic DMEM with ribose for the full 15 days. Day 0 time points were taken from culture at 1 hour.

Riboflavin treated constructs were incubated in 0.25 mM or 0.75 mM riboflavin (Sigma-Aldrich) in PBS for 1 hour after fabrication and then exposed to blue light (BoNew, USA, Dental 10W Cordless LED Light Lamp 2000mw, 420-480 nm) for 40 seconds to trigger non-enzymatic glycation crosslinking.^38^ Blue light was chosen as the photo-activator with a 40 second exposure time due to previous studies showing little impact on chondrocyte cell viability with 0.25 mM and 0.75 mM riboflavin concentration.^38^ After blue light exposure, the constructs were cultured in control DMEM media for the duration of the study. Media was changed every 3 days and constructs were collected for timepoints directly after blue light exposure (day 0) or after 15 days of culture.

Ribose treated constructs were cultured in 30, 100, or 200 mM ribose (Sigma-Aldrich) for the duration of culture. Ribose glycation media consisted of the control DMEM media formulation plus ribose at the desired concentrations. Media was changed every 3 days, and timepoints were taken at 0, 3, 6, 9, 12, and 15 days. Day 0 constructs were soaked in ribose for 1 hour prior to removal from culture to mirror riboflavin incubation periods at Day 0.

### AlamarBlue analysis

AlamarBlue (Bio-Rad) analysis was performed on days 0, 3, 6, 9, 12, and 15 for control and ribose treated constructs and day 0 and 15 for riboflavin-465 nm treated constructs (N = 4) to directly assess metabolism and indirectly assess viability of cells throughout culture. Constructs were placed in 10% alamarBlue and 90% of their respective media formulation. Each media formulation, without constructs, was additionally assessed as controls at each timepoint. The constructs and media controls were kept in the incubator for 7 hours and then media was removed and read using a Biotek Synergy HTX Multi-Mode Reader at wavelengths of 570 nm and 600 nm. Percent reduction of the alamarBlue, indicating active metabolism, was determined from the absorbance readings using the standard manufacturer protocol (Bio-Rad). To account for changes in cell number throughout culture, percent reduction of alamarBlue was also normalized to average DNA/wet weight at each timepoint.

### Post culture analysis

A separate set of constructs, not exposed to alamarBlue, were cultured for up to 15 days (N = 4-7 for each culture condition). At the completion of culture, constructs were photographed to evaluate contraction, and sectioned into pieces that were fixed or frozen for analysis of collagen organization, cellular concentration, collagen content, and fluorescent AGE crosslinks. Additionally, tendons from 21-23 month-old C57BL/6 mouse tails (N=3) were obtained (Jackson Laboratory) for biochemical and fluorescent AGE crosslink analysis to serve as native controls.

#### Collagen fibril analysis

Cross-sections of constructs at day 0 and day 15 were fixed in 10% formalin, stored in 70% ethanol, and imaged with confocal reflectance microscopy to analyze collagen fiber organization as previously described.^45,46,50,51^ Briefly, confocal reflectance imaging was performed in conjunction with fluorescence imaging by splitting a 488 nm laser on a Zeiss LSM 710 Laser Scanning Microscope using a Plan-Apochromat 63x/1.4 Oil DIC M27 (FWD=0.19mm) objective. Confocal reflectance was captured at 400-465 nm by collecting backscattered light reflected by collagen fibers through a 29 μm pinhole and a pixel dwell time of 0.79 μs. Auto-fluorescent cells were captured at 509-577 nm. Noise was reduced by collecting images at 1024 x 1024 pixels with a line average of 4.

Once confocal reflectance analysis was complete, samples were dried with critical point drying, coated with platinum, and imaged with scanning electron microscopy (SEM) to analyze nanometer fibril level diameters. SEM was performed using a Hitachi SU-70-SEM, with samples imaged at a working distance of 15 mm, 5 kV, and 20,000x magnification. Three to five representative images were obtained from N=3 samples per condition. Diameters of 15 fibrils per image were measured with FIJI (NIH) and pooled to determine the average fibril diameter for each sample (total of 45-75 fibrils per samples), then replicate samples from each condition were averaged to determine fibril diameters and standard deviation for each condition.

#### Collagen thermal stability

Thermal stability of collagen was evaluated by differential scanning calorimetry (DSC) as an indirect measurement of overall crosslinking as previously described.^52^ Briefly, frozen day 0 and 15 samples (N = 3-4) were thawed in protease inhibitor for 1 hour. The samples were then drained of excess protease inhibitor on filter paper, weighed (samples ranged from 1-7 mg), and sealed in aluminum pans with 10 μL of PBS. Once sealed, the samples were scanned from 30 to 130°C at a scanning rate of 5°C min^-1^ using a Thermal Advantage Q20 calorimeter (TA Instruments). As a reference an aluminum pan containing 10 μL of PBS was used. The peak temperature of denaturation at maximum heat absorption (T_m_) was determined from the curve using a software integrated with the calorimeter.

#### Biochemical analysis

Biochemical analysis for DNA, GAG, and collagen content was performed as previously reported.^45,46,48,50^ Briefly, half of each construct and mouse tendons were weighted wet (WW), frozen, lyophilized for 48 hours, and weighed dry (DW). Specifically, biopsy punches were removed from culture, cut in half on glass plates, and immediately weighed to determine wet weight. Mouse tails were thawed in PBS, tendons were pulled from the tails, cut in half on glass plates, and immediately weighted to determine wet weight. The samples were then digested at 60°C in 1.25 mg/mL papain solution for 16 hours and analyzed for DNA, GAG, and collagen content via a QuantiFlour dsDNA assay kit (Promega), a modified 1,9-dimethylmethylene blue (Sigma-Aldrich) assay at pH 1.5,^53,54^ and a modified hydroxyproline (hypro) assay,^55^ respectively. Specifically, hydroxyproline content was determined by alkaline hydrolysis in 2N sodium hydroxide for 18 hours at 110°C, samples were then neutralized in hydrochloric acid, oxidized in sodium peroxide, and finally chromophore was produced in 4-(Dimethylamino)benzaldehyde (Sigma-Aldrich) dissolved in n-propanol, and compared to trans-4-Hydroxy-L-proline (Sigma-Aldrich) as standards.^32,45,46,48^ Additionally, construct hydration (percent water) was calculated by determining the difference between the wet weight and dry weight of the sample, and normalizing to the wet weight of the sample.

#### Fluorescent AGE quantification

The same samples used for biochemical analysis were also analyzed for fluorescent AGEs after papain digestion. Fluorometric assays were carried out to assess the effects of ribose and riboflavin-465 nm on the formation of fluorescent AGEs and the AGE crosslink, pentosidine, as previously described.^7,24,29,43,52,56–61^ Briefly, crosslinks were quantified by fluorescence using a Biotek Synergy HTX Multi-Mode Reader at two different ranges of wavelengths. General fluorescent AGEs were quantified by reading fluorescence at 360 nm excitation, 460 nm emission and compared to quinine standards (Sigma-Aldrich) as previously reported.^24,56,60,61^ Pentosidine fluorescence was quantified by reading fluorescence at 328 nm excitation, 378 nm emission^7,52^ and compared to pentosidine standards (Caymen Chemical). Fluorescence at 328 nm/378 nm excitation/emission is consistent with the formation of pentosidine crosslinks;^7,25–27,52,62^ however this range is not specific to pentosidine and may include other auto-fluorescence. Total fluorescent AGEs and crosslinks consistent with pentosidine were normalized to wet weight and hydroxyproline content. The quantity of total fluorescent AGEs and pentosidine crosslinks in constructs were further compared to aged-mouse tail tendons processed similarly to constructs and to reported human pentosidine levels in 60-70 year old menisci^26^ and 80-90 year old cartilage.^25^

### Evaluation of acellular constructs

As a means to separate the effect of cells versus glycation on construct properties, a sub-set of culture conditions were repeated with acellular gels. Collagen gels were created as described above, with the only difference being media was mixed into collagen without cells to adjust to the final concentration of 20 mg/ml. Six mm biopsies were obtained from sheet gels and divided between DMEM control, 0.25 mM riboflavin-465 nm, 0.75 mM riboflavin-465 nm, and 100 mM ribose groups and cultured for up to 15 days identical to cell-seeded constructs, with media changed every 3 days. At the completion of culture, constructs were sectioned into pieces that were fixed or frozen for analysis of collagen fibril organization, collagen content, and fluorescent AGE crosslinks as described above.

### Evaluation of synergistic effect of riboflavin-465 nm and ribose

We also investigated whether ribose and riboflavin-465 nm treatment could be combined to synergistically increase AGE crosslink formation. Ribose and riboflavin-465 treatments, which both significantly increased AGE fluorescence individually, were combined over 15 days of culture to evaluate synergistic effect. Specifically, combination constructs were cultured for 15 days in the 100 mM ribose media with timepoints at day 0, 3, 6, 9, 12, and 15. Prior to removing constructs from culture at each timepoint, constructs were soaked in 0.25 mM riboflavin for 1 hour and induced with blue light for 40 seconds. Each construct was only induced with riboflavin-465 nm treatment once, right before being removed from culture. Constructs were analyzed with the same post-culture analysis described above, specifically evaluating collagen content, total fluorescent AGE accumulation, and pentosidine accumulation.

### Statistics

SPSS was used to confirm normality of data within each group and detect outliers using Shapiro-Wilk tests and Q-Q plots. After confirming normality, all data was analyzed by 2-way ANOVA with Tukey’s post hoc analysis (SigmaPlot 14). All riboflavin treatment analysis was performed separately from ribose treatment analysis as depicted in the separate graphical representation of the data. For DSC analysis, in addition to analyzing similar to all other data, riboflavin and ribose concentrations were pooled at day 0 and day 15 and compared to DMEM constructs by 2 way ANOVA separately. For all tests, *p*<0.05 was threshold for statistical significance. All data are expressed as mean ± standard deviation.

## Results

### Effect of glycation on gross contraction

Inspection of constructs upon removal from culture demonstrated a slight decrease in size in DMEM control constructs and low concentration riboflavin-465 nm (0.25 mM) constructs by day 15, suggesting reduced cellular contraction with increased glycation (0.75 mM riboflavin-465 nm, 30 mM, 100 mM, and 200 mM ribose, **Figure 2A**). However, analysis of construct percent water did not show any significant differences or trends between control and glycated (riboflavin-465 nm and ribose) constructs (**Figure 2B**), indicating negligible differences in degree of contraction between groups.

**Figure 2:**
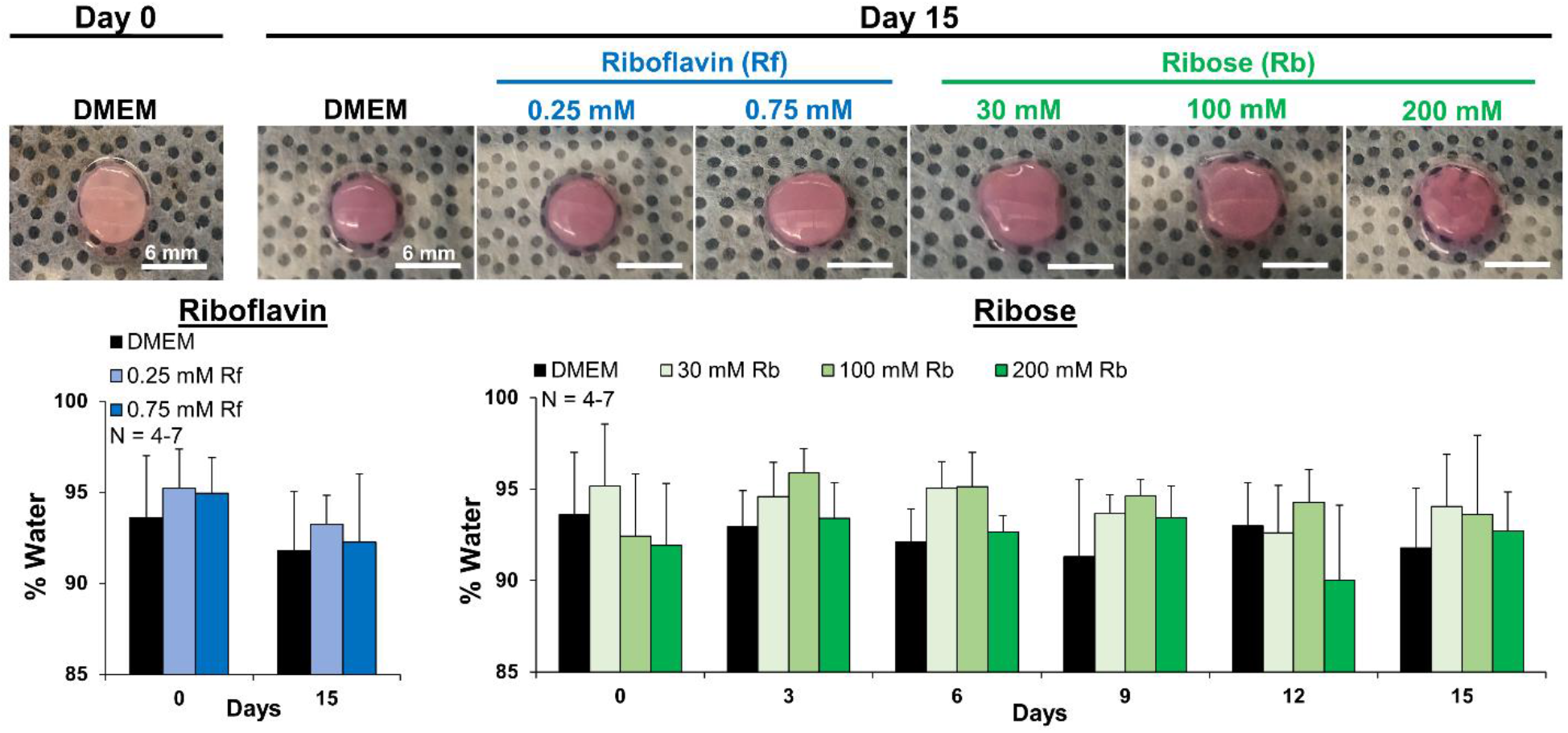
Representative photos of biopsy punches at day 0 and day 15 and percent water concentration of constructs throughout the duration of the study. There were no significant differences in degree of contraction between groups over the 15 days.

### Effect of glycation on collagen fibril organization

SEM and confocal reflectance analysis were used as a macroscale measure to evaluate degree of crosslinking and the effect of AGE induction on collagen fibril organization. Evaluation of collagen organization revealed both glycation systems resulted in increased fibril diameter and organization at the nanometer (SEM) and micrometer (Confocal) length scale by day 15 compared to DMEM controls (**Figure 3**). SEM analysis revealed both concentrations of riboflavin-465 nm treatment significantly increased fibril diameter ~20% compared to DMEM controls at day 0, reaching 39-41 nm diameters, which were maintained through day 15 (**Table 1**). High concentration ribose constructs (100 mM and 200 mM) had a significant increase in fibril diameter compared to DMEM controls at day 0 after 1 hour of culture, and by day 15 all concentrations of ribose resulted in fibril diameters of ~42 nm, a significant 22-23% increase over DMEM controls (**Table 1**). Confocal reflectance revealed DMEM controls had no changes in collagen organization at the micrometer length-scale between day 0 and day 15 (**Figure 3**). However, all concentrations of riboflavin-465 nm and ribose treated constructs developed more organized collagen fibrils by day 15, with more pronounced, elongated fibrils in ribose treated constructs. This clear change in collagen fibril diameter and organization suggests both treatment conditions are crosslinking the collagen, resulting in larger length-scale fibril organiation.^29,33^

**Figure 3:**
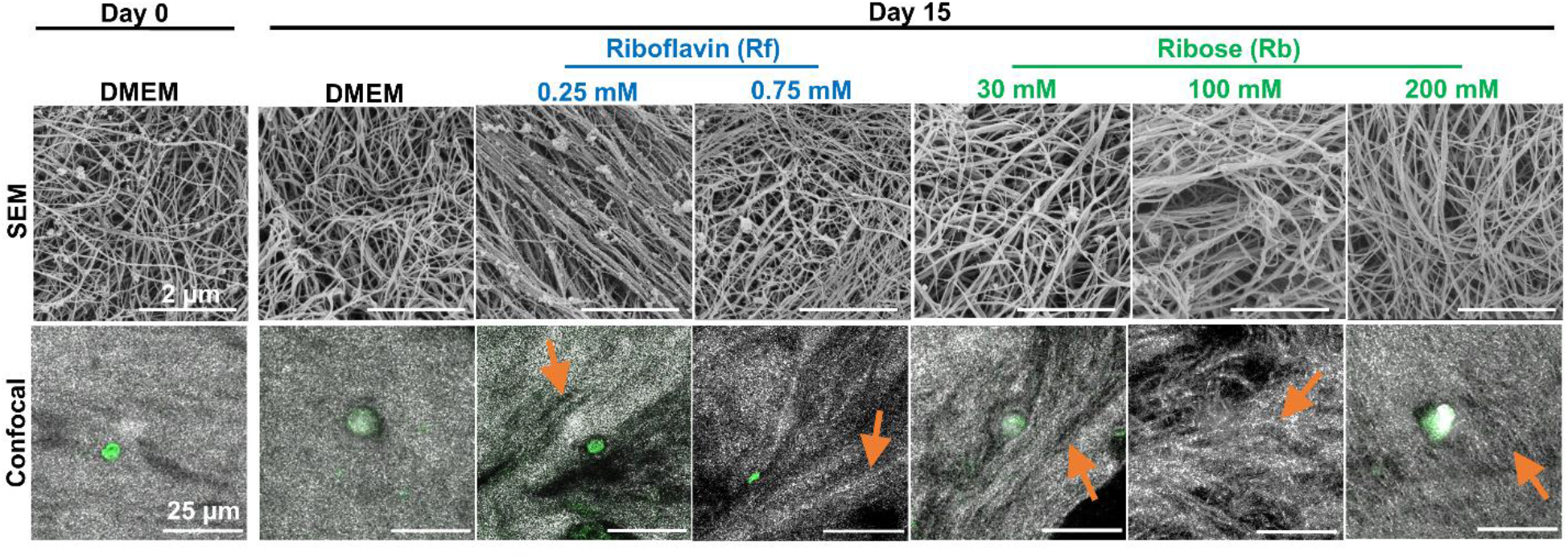
Effect of glycation on collagen organization. SEM analysis revealed similar nanometer diameter fibril organization across all samples, with increasing fibril diameter in riboflavin-465 nm and ribose constructs. Confocal reflectance revealed elongated fibril-like organization at the micrometer length-scale (orange arrows) in all riboflavin-465 nm and ribose constructs compared to DMEM control constructs at day 15. Collagen = grey, auto-fluorescent cells = green.

**Table 1:**
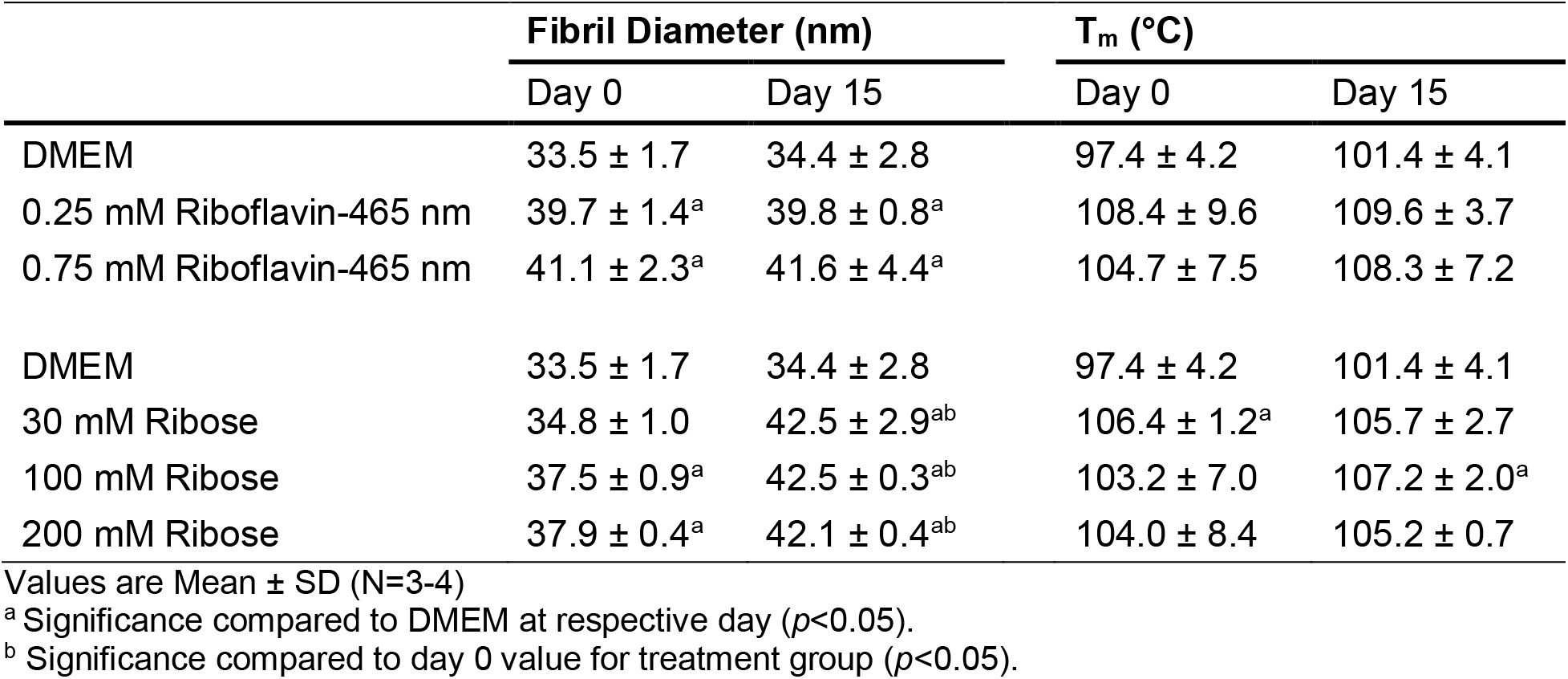
Average fibril diameter and peak temperature of denaturation at maximum heat absorption (T_m_)

| | Fibril Diameter (nm) | | $T_m$ (°C) | |
| --- | --- | --- | --- | --- |
|  | Day 0 | Day 15 | Day 0 | Day 15 |
| DMEM | 33.5 ± 1.7 | 34.4 ± 2.8 | 97.4 ± 4.2 | 101.4 ± 4.1 |
| 0.25 mM Riboflavin-465 nm | 39.7 ± 1.4<sup>a</sup> | 39.8 ± 0.8<sup>a</sup> | 108.4 ± 9.6 | 109.6 ± 3.7 |
| 0.75 mM Riboflavin-465 nm | 41.1 ± 2.3<sup>a</sup> | 41.6 ± 4.4<sup>a</sup> | 104.7 ± 7.5 | 108.3 ± 7.2 |
| DMEM | 33.5 ± 1.7 | 34.4 ± 2.8 | 97.4 ± 4.2 | 101.4 ± 4.1 |
| 30 mM Ribose | 34.8 ± 1.0 | 42.5 ± 2.9<sup>ab</sup> | 106.4 ± 1.2<sup>a</sup> | 105.7 ± 2.7 |
| 100 mM Ribose | 37.5 ± 0.9<sup>a</sup> | 42.5 ± 0.3<sup>ab</sup> | 103.2 ± 7.0 | 107.2 ± 2.0<sup>a</sup> |
| 200 mM Ribose | 37.9 ± 0.4<sup>a</sup> | 42.1 ± 0.4<sup>ab</sup> | 104.0 ± 8.4 | 105.2 ± 0.7 |

Values are Mean ± SD (N=3-4)
<sup>a</sup> Significance compared to DMEM at respective day ( $p < 0.05$ ).<sup>b</sup> Significance compared to day 0 value for treatment group ( $p < 0.05$ ).***Effects of glycation on collagen thermal stability***

Acellular collagen gels had a similar change in collagen fibril organization compared to cellular gels. Both concentrations of riboflavin-465 nm treated acellular gels had a significant 20% increase in fibril diameter at day 0, which was maintained through day 15, while 100 mM ribose treatment significantly increased fibril diameter 23% compared to DMEM controls by day 15 (**Supplemental Table 1**). Further, at the micrometer-length scale both riboflavin-465 nm and 100 mM ribose treatment had increased collagen fibril organization compared to DMEM controls (**Supplemental Figure 1**). This similar change in collagen fibril diameter and organization with and without cells further suggests the changes in collagen organization are from glycation induced crosslinking and not primarily cellular induced changes.

### Effects of glycation on collagen thermal stability

Both riboflavin-465 nm and ribose treatment increased the denaturation temperature (T_m_) of collagen gels with time in culture, suggesting both glycation techniques induce crosslinking.^52,63^ Although conditions were not significant individually, pooling both concentrations of riboflavin-465 nm treatment resulted in a significant increase in the denaturation temperature from 97.4 ±4.1°C (day 0 DMEM construct T_m_) to an average 106.5 ±8.1°C after blue light treatment at day 0, and they maintained a significantly higher T_m_ of 108.9 ±5.6°C compared to DMEM controls at day 15 (**Table 1**). Similarly, pooled ribose treated constructs had a significant increase in denaturation temperature compared to DMEM control constructs to 104.6 ±5.9°C at day 0 and 106.2 ±2.0°C at day 15 (**Table 1**).

### Effects of glycation on cellular viability and metabolism

To evaluate the effect of riboflavin-465 nm and ribose on cell viability and metabolism, construct DNA was measured as a representation of cell number, and reduction of alamarBlue was measured as a representation of metabolic activity. DNA analysis demonstrated both concentrations of riboflavin-465 nm (0.25 mM and 0.75 mM) had no change in DNA concentration compared to control at day 0 (**Figure 4A**). Further, both concentrations of riboflavin-465 nm treated constructs maintained similar DNA to control constructs by day 15, suggesting no change in cell number between treatment groups. Similarly, both concentrations of riboflavin-465 nm treated constructs reduced alamarBlue similar to control constructs at day 0 and day 15 when normalized to DNA concentration, suggesting little to no effect on cellular metabolism. (**Figure 4B**). Collectively this suggests both concentrations of riboflavin-465 nm treatment had little effect on cell viability and metabolism. While 0.75 mM riboflavin-465 nm constructs did not have a different DNA concentration compared to 0.25 mM riboflavin-465 nm and control constructs at day 15, they did lack an increase in DNA compared to day 0, which could suggest a decrease in cell number or proliferation. Previously, it has been shown that higher concentrations of riboflavin lead to reduced long-term viability.^38^ This is believed to be due to increased free radical formation during irradiation with blue light.^38^ However, 0.75 mM riboflavin-465 nm constructs did significantly increase reduction of alamarBlue at day 15, so cells may have shifted to being more metabolically active and less proliferative.

**Figure 4:**
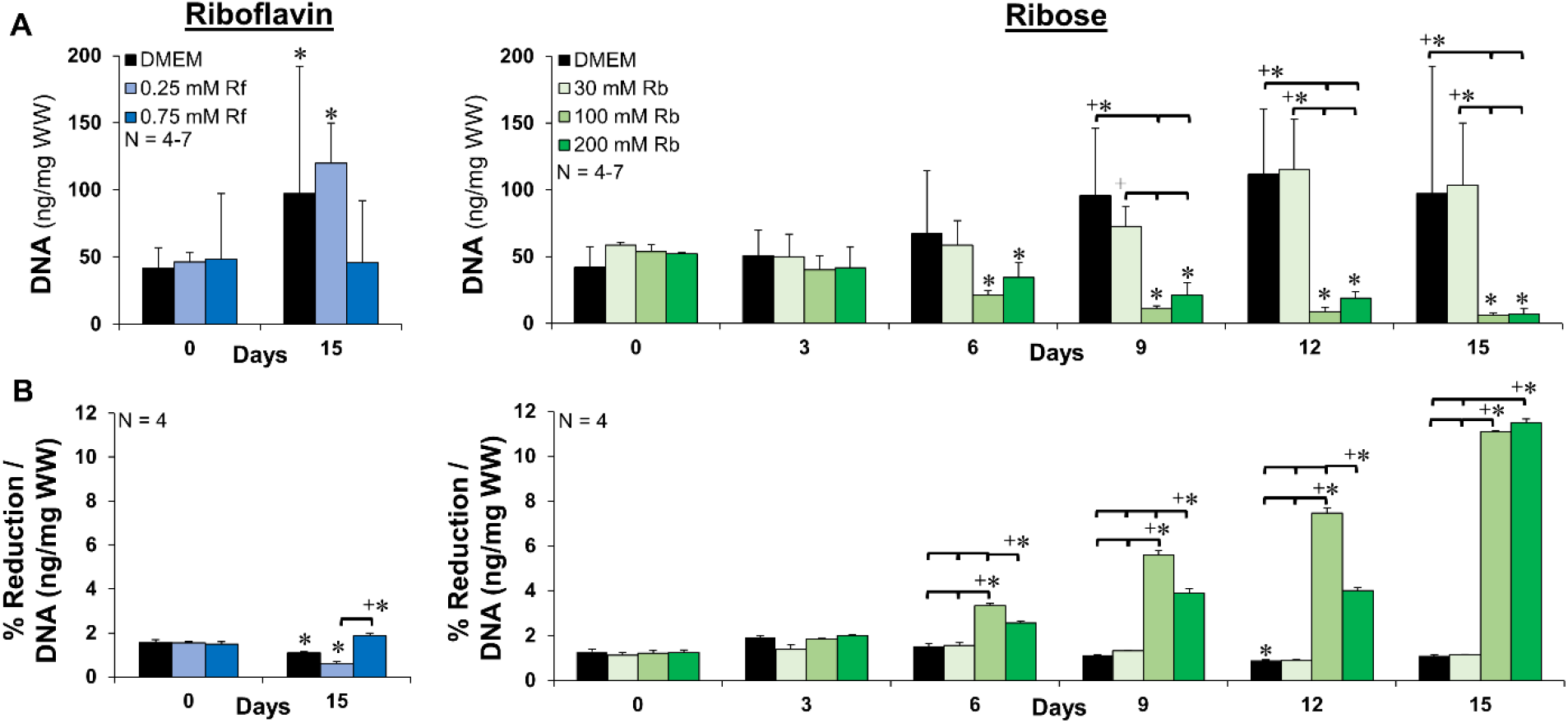
Effect of glycation on cell count and metabolism. A) DNA normalized to wet weight (WW) and B) alamarBlue percent reduction normalized to average DNA/WW. Riboflavin-465 nm had little effect on cell number and metabolism, while higher concentrations of ribose significantly decreased cell number starting at day 6 and significantly increased cellular metabolism by day 6. Significance compared to *day 0 of treatment group or +bracketed group (*p*<0.05).

In contrast, high concentrations of ribose (100 mM and 200 mM) produced significant decreases in DNA by day 6, while 30 mM ribose treatment maintained similar DNA to control constructs throughout culture (**Figure 4A**). Further, both high concentrations of ribose (100 mM and 200 mM) resulted in significantly lower reduction of alamarBlue compared to control and 30 mM ribose constructs (**Supplemental Figure 2**); however, when normalized to DNA concentration, both high concentrations of ribose had significantly higher reduction of alamarBlue starting at day 6 (**Figure 4B**). Collectively this suggests high concentrations of ribose reduce cell viability while increasing cellular metabolism with time in culture.

Glycosaminoglycan (GAG) accumulation, normalized to DNA, was quantified as an additional measure of metabolism since this is a major macromolecule often produced by meniscal fibrochondrocytes (**Supplemental Figure 2**). Similar to alamarBlue analysis, both concentrations of riboflavin-465 nm had little change in GAG accumulation compared to control constructs throughout culture, while high concentrations of ribose (100 mM and 200 mM) resulted in significantly higher GAG accumulation by day 15.

### Induced Fluorescent AGE Crosslink Accumulation

In this study, total fluorescent AGEs and AGE crosslink pentosidine, were measured to assess the ability of riboflavin-465 nm and ribose to induce AGEs. Collagen concentration, represented by hydroxyproline (hypro) content, was measured to normalize AGE crosslinks across all treatment groups. As expected, collagen content was relatively constant throughout culture with no consistent differences across DMEM, riboflavin-465 nm, and ribose groups (**Figure 5A**).

**Figure 5:**
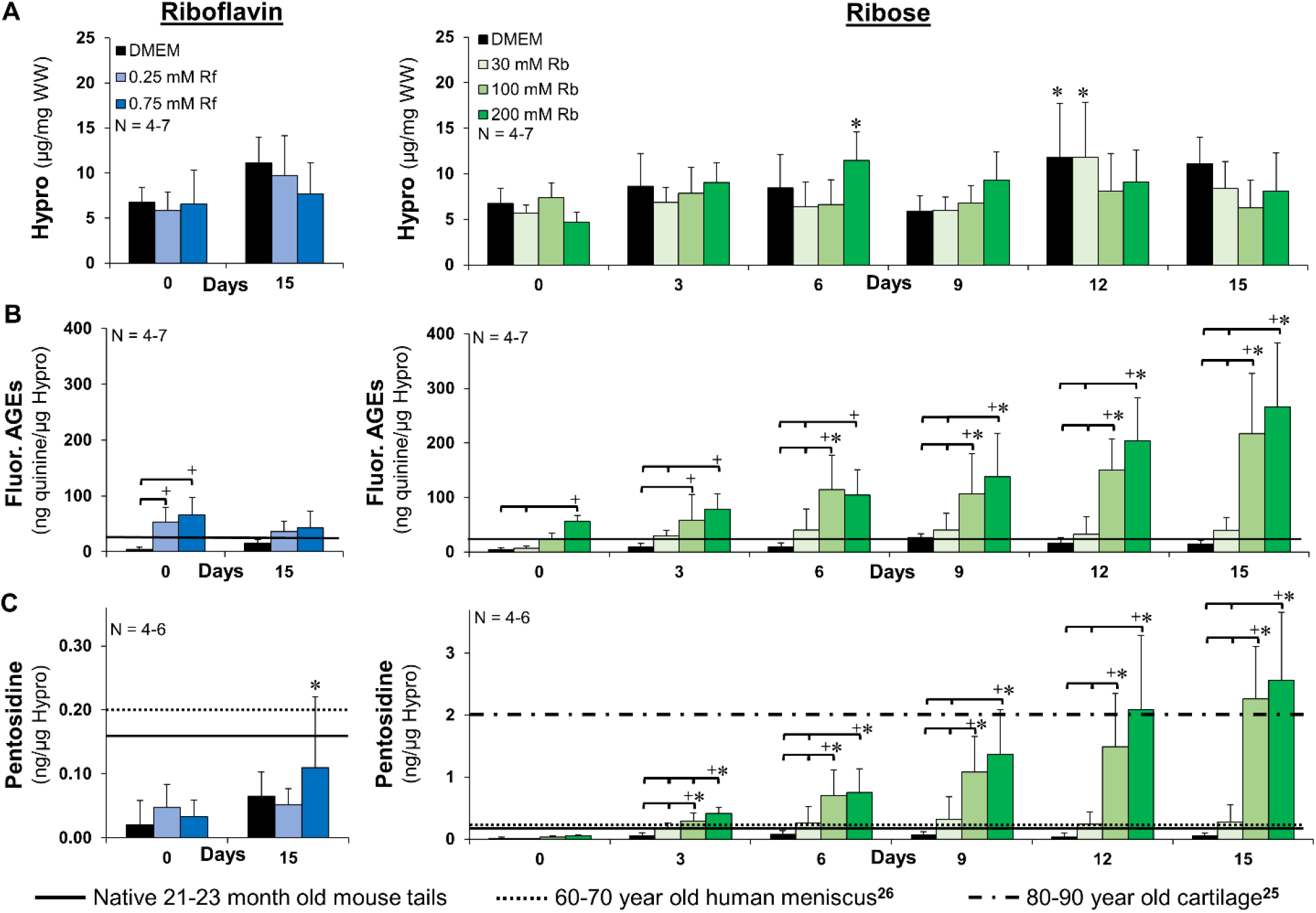
Glycation significantly increases AGE formation. A) Collagen content represented by hydroxyproline (Hypro) normalized to wet weight (WW), B) fluorescent AGEs and C) AGE crosslink pentosidine determined by pentosidine auto-fluorescence, normalized to hypro. Both concentrations of riboflavin-465 nm induced native levels of fluorescent AGEs, but had little effect on pentosidine accumulation. Ribose induced fluorescent AGEs and pentosidine crosslinks in a dose dependent fashion, reaching aged C57BL/6 mouse and human values^25,26^ between 3 and 15 days. Significance compared to *day 0 of treatment group or +bracketed group (*p*<0.05).

Riboflavin-465 nm and ribose induced fluorescent AGEs in a dose dependent fashion (**Figure 5**). Specifically, both concentrations of riboflavin-465 nm induced significant increases in fluorescent AGE accumulation normalized to wet weight (**Supplemental Figure 3A**) and collagen content (**Figure 5B**) at day 0 and maintained a similar level of fluorescent AGEs through 15 days of culture. Further, both concentrations of riboflavin-465 nm treatment exceeded aged 21-23 month old mouse tail tendon fluorescent AGE levels after induction at day 0 (**Figure 5B**, solid line). Ribose treatment significantly increased fluorescent AGEs when normalized to wet weight (**Supplemental Figure 3A**) and collagen content (**Figure 5B**) in a dose dependent fashion with time in culture. Further, all concentrations of ribose matched or exceeded aged mouse tail tendon fluorescent AGE levels by 3 days of culture (**Figure 5B**).

Despite riboflavin-465 nm significantly increasing fluorescent AGEs, there were no significant increases in fluorescent pentosidine levels compared to DMEM controls, when normalized to wet weight (**Supplemental Figure 3B**) or collagen content (**Figure 5C**). However, 0.75 mM riboflavin-465 nm constructs did have a significant increase in pentosidine by day 15, compared to day 0. Further, riboflavin-465 nm treatment did not match pentosidine levels of aged mouse tail tendon or aged human meniscus.^26^ Alternatively, ribose treatment did significantly increase pentosidine crosslinks when normalized to wet weight (**Supplemental Figure 3B**) and collagen content (**Figure 5C**) in a dose dependent fashion, similar to fluorescent AGE accumulation. Further, 100 mM and 200 mM ribose induced pentosidine concentrations that matched aged mouse tail tendon and reported 60-70 year old human meniscus pentosidine levels by 3 days.^26^ Additionally, 80-90 year old human cartilage pentosidine levels^25^ were reached by day 12 with 200 mM ribose treatment, and by day 15 with 100 mM ribose treatment (**Figure 5**).

Acellular collagen gels had no significant difference in fluorescent AGE accumulation compared to cell-seeded collagen gels throughout culture for both riboflavin-465 nm (0.25 mM and 0.75 mM) and 100 mM ribose treatment (**Supplemental Figure 4A**). Similarly, acellular and cell-seeded collagen gels had no significant difference in pentosidine accumulation for both riboflavin-465 nm concentrations; however, interestingly cell-seeded constructs treated with 100 mM ribose did have reduced pentosidine accumulation compared to 100 mM ribose acellular constructs from day 9 on (**Supplemental Figure 4B**). Pentosidine crosslinks take longer to form than many other AGEs,^64^ thus it is possible cellular turn-over or interactions with the matrix are reducing the formation of pentosidine in ribose treated constructs.

### Synergistic Effect of Riboflavin-465 nm and Ribose Treatment

In this study we also investigated whether ribose and riboflavin-465 nm treatment could be combined to synergistically increase AGE crosslink formation, making it possible to reach 80-90 year old aged cartilage levels earlier in culture before the ribose induced viability issues. Combination constructs were cultured for 15 days in 100 mM ribose media and prior to removing constructs from culture at each timepoint, constructs were soaked in 0.25 mM riboflavin and induced with blue light for 40 seconds. Surprisingly, we found there was no synergistic increases in crosslinks when riboflavin-465 nm and ribose were combined at any of the timepoints. The combination constructs had the same amount of fluorescent AGEs as riboflavin-465 nm treatment alone at day 0 and did not further significantly increase the amount of fluorescent AGEs until day 12. Further, the combination constructs had similar concentrations of fluorescent AGEs and pentosidine crosslinks as the 100 mM ribose treated constructs throughout culture (**Figure 6**).

**Figure 6:**
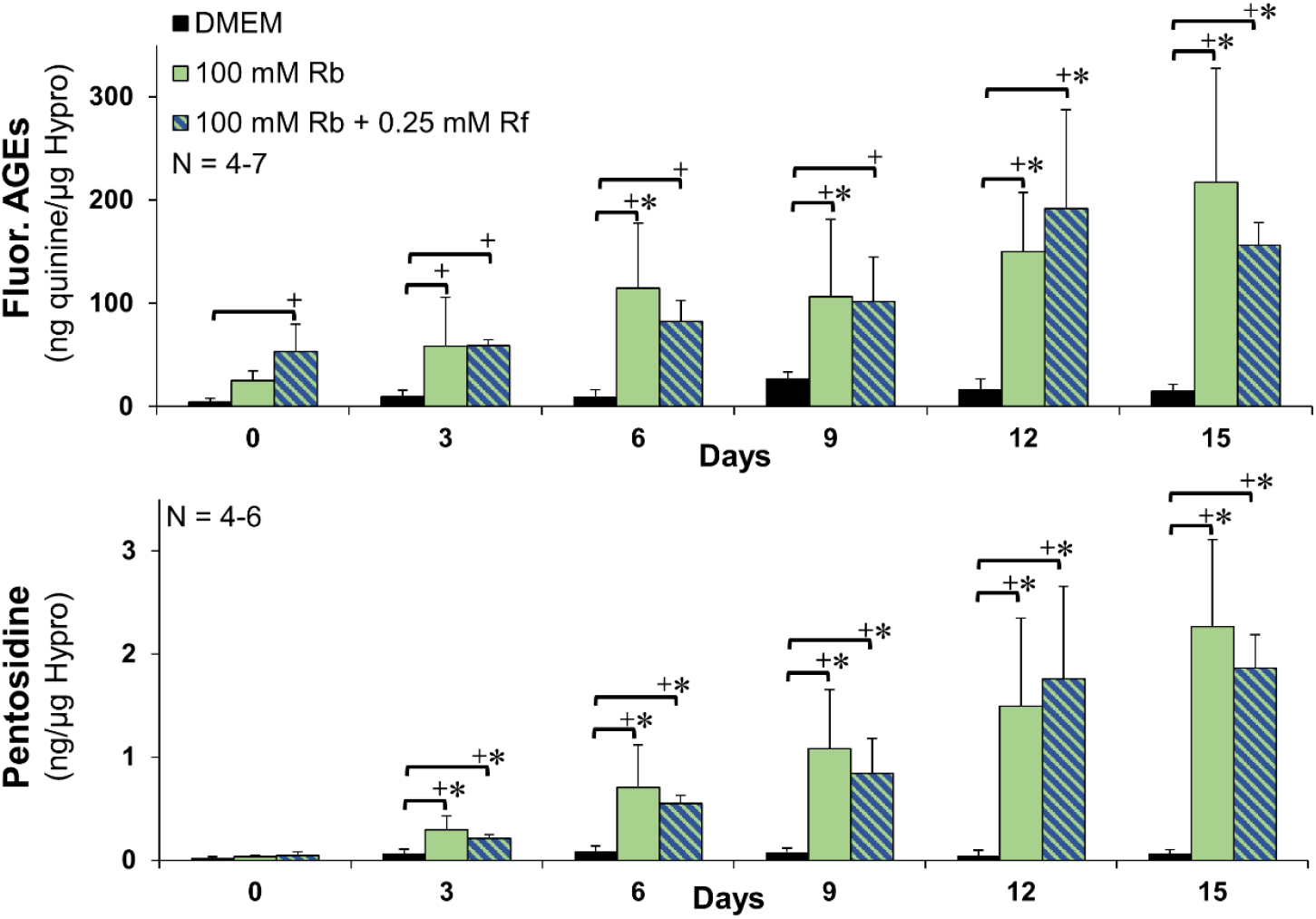
Combination system does not synergistically increase fluorescent AGE or pentosidine formation. Fluorescent AGE and AGE crosslink pentosidine, normalized to hydroxyproline (hypro). 100 mM ribose (Rb) alone and in combination with 0.25 mM riboflavin-465 nm (Rf) induced fluorescent AGEs and pentosidine crosslinks in a similar dose dependent fashion. Significance compared to *day 0 of treatment group or + bracketed group (*p*<0.05).

## Discussion

It has been projected that the number of people over 65 years of age in 2015 will triple by 2050, making up approximately 17% of the worldwide population.^65^ This increase in the amount of people living past age 65, suggests a better understanding of the link between aging and tissue degeneration could have major treatment and socioeconomic impacts.^66^ Despite ample correlative evidence connecting AGEs to tissue injury,^63,67^ degeneration,^24,27^ and multiple chronic diseases, such as cardiovascular disease,^18^ chronic kidney disease,^19^ diabetes,^68^ Alzheimer’s and Parkinson’s disease,^23^ there are many un-answered questions related to AGEs impact on cell-matrix interactions, resulting in limited therapeutic options. One reason for this limited understanding is because AGEs are typically induced using concentrations of ribose greater than 200 mM, which decrease cell viability and make it impossible to investigate cell-matrix interactions.^7,10,29^ Here, we induced native levels of fluorescent AGEs with lower concentrations of ribose and maintained cell viability through 15 days of culture. Specifically, in this study we found ribose and riboflavin-465 nm are capable of producing fluorescent AGE crosslinks which match and/or exceed reported human fluorescent AGE levels for various connective tissues, ages, and diseases.^7,24–27^ Interestingly, a single 40 second treatment of 0.25 mM or 0.75 mM riboflavin and blue light produced similar levels of fluorescent AGEs as 1 hour of 200 mM ribose treatment and 3 days of 100 mM ribose treatment, without decreasing viability. This riboflavin-465 nm treatment option is a promising means to trigger AGE crosslinks on demand *in vivo* or *in vitro* without impacting cell metabolism with prolonged cultures in high ribose concentrations.

There are several different AGEs that occur throughout the body. The mechanism by which they individually form remains unclear, however this population of AGEs includes adducts which modify the surface of a protein and crosslinks that bridge two neighboring proteins.^9,69^ In this study, we evaluated both total fluorescent AGE accumulation and AGE crosslink pentosidine accumulation. AGEs are typically categorized into four different types; fluorescent crosslinked or uncrosslinked, and non-fluorescent crosslinked or uncrosslinked.^15^ Here we measured total fluorescent AGE accumulation using previously established fluorescence measurements at 360 nm excitation and 460 nm emission.^24,29,43,52,56–61^ These specific wavelengths are consistent with all AGEs that naturally fluoresce, including pentosidine, pentodilysine, crossline, pyrropyridine, Vespertysine A, Vespertysine C, and fluorescent AGE adducts.^15^ In addition to measuring fluorescent AGE accumulation we also measured AGE crosslink pentosidine auto-fluorescence. While pentosidine is not the most common AGE, it is a major type of AGE which is commonly studied due to its easily identified fluorescent wavelength found to be consistent with its formation,^62^ making it easy to analyze using fluorescence measures.^7,22,25–27,52^ Pentosidine has been found in high levels in patients with rheumatoid arthritis,^21,22^ plays a pivotal role in the complications of diabetes,^21,70^ and has even been linked to being a risk factor for patients with heart failure.^71^ While pentosidine only represents a small portion of total AGEs, it is an important AGE crosslink and being able to induce human pentosidine levels without decreasing cell viability will allow for pentosidine and its effects to be studied in a way that has not been done before in *in vitro* models.

Both the ribose and riboflavin-465 nm glycation methods resulted in increased collagen fibril diameters and increased thermal stability by 15 days suggesting induced crosslinking.^29,33^ Collagen gels produced by acid extraction techniques, as was performed in this study, self-assemble into fibrils with diameters of 20-70 nm within a few hours of gelation.^72^ Altering the pH, collagen concentration, ion concentration, or temperature of gelation will alter the size of fibrils formed.^72,73^ As observed by SEM, our collagen gels form collagen fibrils ~33 nm in diameter when first gelled. Riboflavin-465 nm constructs had a significant 20% increase in fibril diameter right after treatment on day 0, while ribose treated constructs had a significant 23% increase in fibril size by day 15 (**Table 1**). Interestingly, while acellular gels had smaller fibril diameters (~29-30 nm), possibly due to less crowding during gelation, they had a similar 20% increase in fibril diameter with riboflavin-465 nm and ribose treatment (**Supplemental Table 1**). Further, when we evaluated collagen fibril organization at a lower magnification with confocal reflectance, we were un-able to observe any collagen organization in DMEM control constructs, and increasingly more fibril-like structures with increasing concentrations of riboflavin-465 nm and ribose treatment (**Figure 3**), suggesting enhanced collagen organization into larger and/or longer collagen fibrils with glycation.

Previously it has been reported that riboflavin treatment, with UV or blue light, and ribose treatment results in significant increases in collagen fibril diameter both at the nanometer scale and micrometer length-scale.^29,33,39,44,74–76^ It is believed glycations increase fibril size on the nanometer scale by inducing crosslinks that push the collagen molecules apart, or pull more water into the fibril, resulting in an increase in intermolecular spacing and diameter of the collagen fibrils,^33,74,75,77^ while on the larger micrometer-level scale it is believed crosslinking increases fibril size due to actively crosslinking more collagen fibrils together.^29,33^ While it is plausible these changes in collagen fibril organization are due to cellular contraction and response to differences in mechanics, the fact acellular gels had a similar 20% increase in fibril diameter and degrees of micrometer organization, suggests these changes are primarily due to crosslinking.

AGEs can form adducts or crosslinks, and while these glycation systems most likely induce both, similar to the *in vivo* environment, the increase in fibril diameter and increase in collagen denaturation temperature suggests we are indeed achieving crosslinking of the collagen fibrils.^52,63^ AGE crosslinking of collagen will not only result in altered mechanics,^7,10^ but can also lead to dramatic modifications of collagen interactions with other molecules such as proteoglycans, enzymes, and cell integrins.^9^ These modifications to the collagen fibril surface are known to affect cell-matrix interactions in a manner that leads to impaired wound healing and increased inflammation.^78^ Our high density collagen system provides a promising means to further investigate these direct effects of AGE crosslinks on collagen to better understand their role in AGE driven connective tissue disease and damage throughout the body.

In this study, the ribose glycation system induced fluorescent AGEs and pentosidine crosslinks in a dose dependent fashion over 15 days, matching reported human fluorescent AGE levels for various tissues, ages, and diseases.^25,26^ Specifically, all concentrations of ribose treatment induced fluorescent AGE accumulation that matched 21-23 month old mouse tail tendons (~similar to 55-65 year old human tissue^79^) by 3 days. Further, ribose induced pentosidine accumulation that matched reported human 60-70 year-old meniscus^26^ by 3 days and 80-90 year-old cartilage by 12 days.^25^ Previous work in the field has induced AGEs using ribose concentrations greater than 200 mM, often resulting in super-physiological levels of glycation and low cell viability, most likely due to hyperosmotic conditions,^7,10,29,32^ making it impossible to investigate cell-matrix interactions. Here, we induced native levels of AGEs with a lower concentration of 30 mM ribose and maintained cell viability through 15 days of culture. As expected, we did observe decreased cell viability with higher concentrations of ribose. Specifically, 100 mM and 200 mM ribose treatments resulted in significantly lower DNA concentrations by day 6, most likely due to a hyperosmotic stress environment.^29,32^ Further, these high ribose concentrations increased cellular metabolism, as evidenced by significantly higher reduction of alamarBlue, a measure of metabolic activity, by day 6 (**Figure 4**) and significantly higher accumulation of GAGs, a major macromolecular produced by meniscal fibrochondrocytes, by day 15 (**Supplemental Figure 1**). Collectively this demonstrates the limitations of inducing AGEs with high concentrations of ribose and why lower concentrations should be considered for evaluating cellular response to AGE crosslinks. In this study, we matched aged mouse tail tendon and human meniscus AGE levels by 3 days, before any changes in cell viability or metabolism.

In an attempt to overcome the cell viability and metabolic changes that occur with ribose treatment, we also investigated the potential of riboflavin-465 nm to induced AGEs. Riboflavin, or B2 vitamin, is a non-toxic photosensitizer that when exposed to UV light or blue light (465 nm) results in the formation of non-enzymatic covalent crosslinks which form similar to AGEs in collagen.^33^ Riboflavin is used clinically to enhance corneal strength and *in vitro* to increase the strength of different hydrogels.^33,36–38,41^ Specifically, it has been shown to crosslink collagen gels in a dose dependent fashion with limited viability issues at the concentrations used in this study.^37,40^ Riboflavin holds great potential to create an inducible AGE model, however despite its wide use for collagen crosslinking, it has not been investigated as a means to induce AGEs. In this study, we found that a single treatment of riboflavin and 40 seconds of blue light produced similar levels of fluorescent AGEs as 1 hour of hyperosmotic 200 mM ribose treatment or 3 days of 30-200 mM ribose treatment, without decreasing viability. Further, this single 40 second treatment of riboflavin-465 nm induced fluorescent AGE accumulation which matched aged mouse tail tendon. The 0.25 mM riboflavin-465 nm treatment had no change in cell viability or metabolism throughout 15 days of culture as well. Previously, it has been suggested that riboflavin, initiated with UV or blue light, crosslinks collagen in a similar manner as AGEs ^33^ and it has been shown AGE inhibitors can block the crosslinks formed by riboflavin.^35^ However, this is the first study to our knowledge to demonstrate that riboflavin-465 nm photo-initiated crosslinking results in AGE crosslink formation. This riboflavin-465 nm treatment option is a promising means to trigger AGE crosslinks on demand *in vivo* or *in vitro* without impacting cell metabolism or cell viability to better study the mechanism of AGEs in connective tissues.

In this study we also investigated whether ribose and riboflavin-465 nm treatment could be combined to have a synergistic increase in fluorescent AGE crosslink formation making it possible to reach 80-90 year old cartilage fluorescent AGE levels earlier in culture, before the ribose induced viability issues. Surprisingly, the results showed that there was no synergistic increase in crosslinks when riboflavin-465 nm and ribose were combined. The combination of riboflavin and ribose produced similar levels of fluorescent AGEs as riboflavin-465 nm treatment alone through 9 days and maintained similar levels of fluorescent AGEs as 100 mM ribose treatment alone throughout the rest of culture (**Figure 6**). This suggests that ribose and riboflavin-465 nm may target similar amino acids in collagen. While it is still not completely understood the specific sites of AGEs in collagen, previously it has been suggested that glycations from ribose that form AGEs primarily target lysine, arginine, and histidine.^35,80–82^ While photo-initiated riboflavin can react with lysine and arginine, these amine groups have been shown to play a smaller role in riboflavin crosslinking.^83^ Instead, riboflavin crosslinking has been suggested to be more dependent on carbonyl groups, such as hydroxyproline, hydroxylysine, tyrosine, and threonine.^83^ However, recent work has demonstrated that ribose can also target hydroxylysine, as well as lysine, arginine, and histidine.^81^ Further, it has been suggested reactive oxygen species (ROS) produced during photo-initiation of riboflavin primarily target histidine.^83^ This combined with the fact that the presence of AGEs can lead to the production of ROS which can then react with amino acids to form more AGEs,^11,12,17,35^ demonstrates that while ribose and riboflavin-465 nm may favor different amino acids, there is a large amount of overlap. Since ribose and riboflavin most likely target similar amino acids of collagen to form AGEs, addition of riboflavin after ribose incubation would not further increase AGE formation if most reactionary amino acids are occupied.

While riboflavin-465 nm treatment induced native mouse fluorescent AGE levels, riboflavin-465 nm treatment did not appear to induce pentosidine crosslinks, as measured by pentosidine auto-fluorescence (**Figure 5**). This is in contrast to what was seen with the ribose glycation method. Pentosidine is formed from sugars reacting with lysine and arginine residues of proteins.^62,64,82,84^ As discussed previously, while photo-initiated riboflavin can react with lysine and arginine, these amino acids play a minor role in riboflavin-initiated crosslinking^83^ and thus it makes sense riboflavin-465 nm would not favor the formation of pentosidine at day 0. Further, pentosidine crosslinks take longer to form than many other AGEs,^64,82,84^ thus it would not be expected to form after 1 hour of treatment, as is also observed in the ribose constructs. Originally, pentosidine was hypothesized to form by reaction of pentose sugars only;^62^ however later studies demonstrated glucose, fructose, ascorbate, tetrose, triose, and other glycated proteins were also possible precursors with longer incubations.^64,82,84^ While pentoses are the most efficient source of pentosidine, these other sugars and glycated proteins can form pentosidine but at slower rates.^82,84^ Thus while riboflavin-465 nm treatment does not appear to significantly form pentosidine at initiation, it is possible it could eventually lead to pentosidine formation, as is seen with 0.75 mM riboflavin-465 nm constructs having a significant increase in pentosidine fluorescence by 15 days, albeit at a much lower concentration than that induced by ribose. Collectively, this data suggests that these two glycation systems can induce different types of AGE crosslinks and opens the door to exploring effects of tissue specific crosslink effects throughout the body.

While the inducible systems investigated in this study are promising for investigating the mechanism of AGEs in the presence of viable cells, there are several limitations that must be considered. One limitation of this study is that we only evaluated AGE induction in collagen since this is the main structural protein of tissues throughout the body and the most common protein evaluated for AGE accumulation. However, other long-lasting proteins in the ECM, such as elastin and fibronectin, are also susceptible to AGEs.^16^ The amount of AGEs induced may vary with different concentrations of collagen gels, different degrees of collagen organization, different proteins in the ECM, or in different disease conditions. Future studies in whole tissue or other systems may have to vary concentration and duration of exposure to reach physiological levels of AGEs; however this work can serve as a guide for inducing physiological levels of fluorescent AGE and pentosidine levels with no change in cell viability or metabolism. Additionally, we only measured AGE accumulation through fluorescence signatures, this neglects all non-fluorescent AGEs that may be formed by these induction systems and does not allow for the distinction between adducts and crosslinks. Further the fluorescence signature used to quantify pentosidine is not specific to pentosidine and may include other auto-fluorescence. Further studies should use high performance liquid chromatography (HPLC) to better distinguish the population of AGEs produced.

Another limitation of this study is that AGEs were induced quickly *in vitro* which does not entirely replicate the *in vivo* process of life-long accumulation of AGEs. This acute induction of AGEs *in vitro* does not allow for cell-mediated matrix adaptations or inflammatory signaling which can further compound the effects of AGEs in age-related disease.^12,14,17^ However, this study does take a significant step towards better mimicking the *in vivo* environment by characterizing systems that induce physiological levels of fluorescent AGEs while maintaining cell viability, providing a means to induce AGEs *in vivo* or in native tissue and then explore cell-matrix interactions without other confounding factors that occur during natural aging.

Similarly, a limitation of inducing AGEs with photo-initiated riboflavin is the production of ROS which can further crosslink collagen and alter cell viability, creating an environment that cells may not encounter *in vivo* during lifetime accumulation of AGEs. As discussed in the introduction, AGEs are believed to induce chronic diseases due to two main mechanisms. The first mechanism is the traditional crosslinking of proteins, altering the structure and mechanics of the tissue. The second mechanism is from AGEs interacting with cellular receptors, primarily RAGE receptor, leading to increased production of ROS and inflammatory cytokines, which in turn can create more AGEs.^12,14,17^ Thus ROS produced from photo-activation of riboflavin may produce changes in the tissues that are similar to those seen in chronic disease with AGEs. In fact, riboflavin crosslinking of corneas using UV light has been shown to produce changes very similar to those seen in aged corneas.^35,80^ Further, it has been shown that AGE inhibitors used during riboflavin-UV treatment block changes in mechanics and collagen crosslinking, suggesting crosslinks formed by ROS are either minor or are AGEs.^35^ However, the production of ROS from riboflavin-465 nm may be quicker and at a much higher concentration than what is produced *in vivo*. This is a concern with photo-activation, particularly for cell viability, thus we have taken steps to reduce the amount of ROS produced by using blue light instead of UV and short exposure times. Blue light has longer wavelengths than UV light, and has been shown to result in less cytotoxicity and less free radicals than UV light treatment.^38,85,86^

## Conclusions

There is increasing evidence that AGEs play a central role in aging and degeneration of connective tissues; however, little is known about how AGEs impact tissue mechanics and damage cell-matrix interactions. Even less is known about how these factors interact to cumulatively affect local repair of matrix damage, inflammation, disease progression, and systemic health. In this study we developed and characterized two inducible AGE models, low concentration ribose and riboflavin with blue light, which produced fluorescent AGE quantities which matched native tissue fluorescence levels without inducing cell viability and metabolic changes. These systems provide promising means to investigate the mechanism of AGEs in altering cell-matrix interactions and matrix mechanics *in vitro* and *in vivo* so to better mitigate the role of AGEs in tissue injury and chronic disease with age and diabetes.

## Acknowledgements

The authors would like to thank are Dr. Rebecca Heise, Dr. Michael Valentine, Dr. Carl Sayer, Dr. Barbara Boyan, and Cydney Dennis for their support in this research. The authors acknowledge the use of facilities within the Nanomaterials Characterization Core at Virginia Commonwealth University.

