## Supplemental for "An Inducible Model for Unraveling the Effects of Advanced Glycation End-Product Accumulation in Aging Connective Tissues"

**Supplemental Table 1:**
Average fibril diameter for acellular gels at day 0 and day 15.

| | Fibril Diameter (nm) | |
|---|---|---|
| | Day 0 | Day 15 |
| DMEM | 29.6 ± 1.2 | 29.5 ± 2.0 |
| 0.25 mM Riboflavin | 35.4 ± 1.6[a] | 34.8 ± 0.5[a] |
| 0.75 mM Riboflavin | 35.3 ± 0.9[a] | 34.6 ± 1.9[a] |
| 100 mM Ribose | 32.5 ± 1.0[a] | 36.5 ± 1.1[ab] |

Values are Mean ± SD (average of 5 images per sample with 15 fibrils measured per image)
[a] Significance compared to DMEM at respective day ($p$<0.05).
[b] Significance compared to day 0 value for treatment group ($p$<0.05).

**Supplemental Figures**

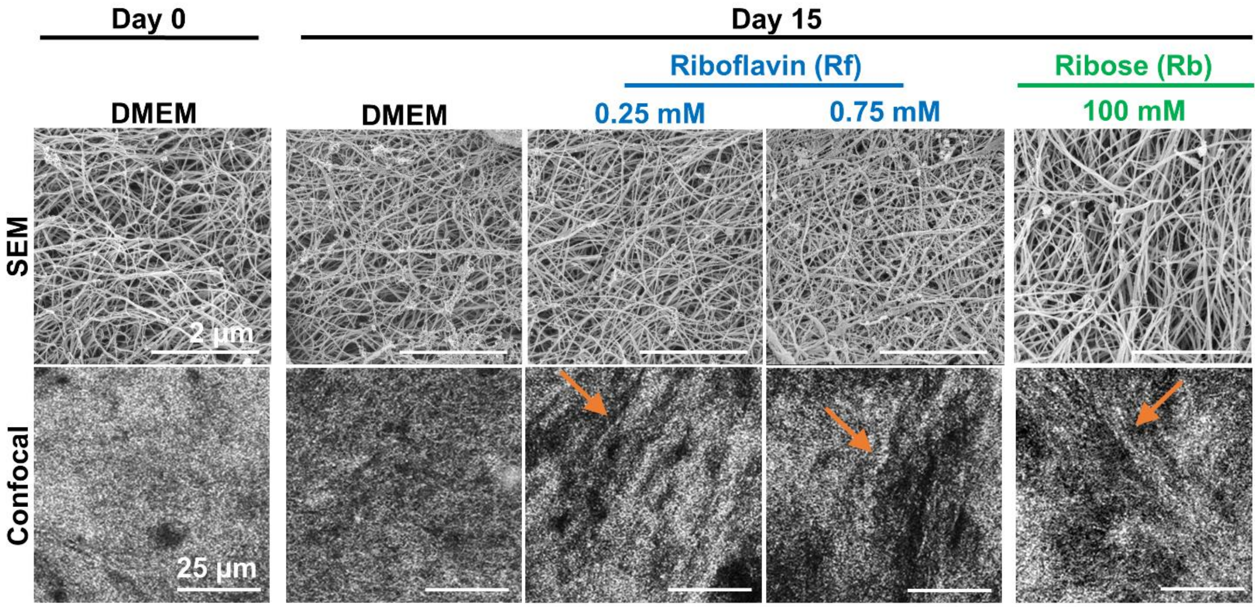


**Supplemental Figure 1:** Effect of glycation on collagen organization in acellular gels. SEM analysis revealed similar nanometer diameter fibril organization across all samples, with increasing fibril diameter in riboflavin-465 nm and ribose acellular gels, similar to cell-seeded constructs, suggesting increased crosslinking. Confocal reflectance revealed elongated fibril-like organization at the micrometer length-scale (orange arrows) in all riboflavin-465 nm and ribose acellular gels compared to DMEM control gels, similar to cell-seeded constructs.

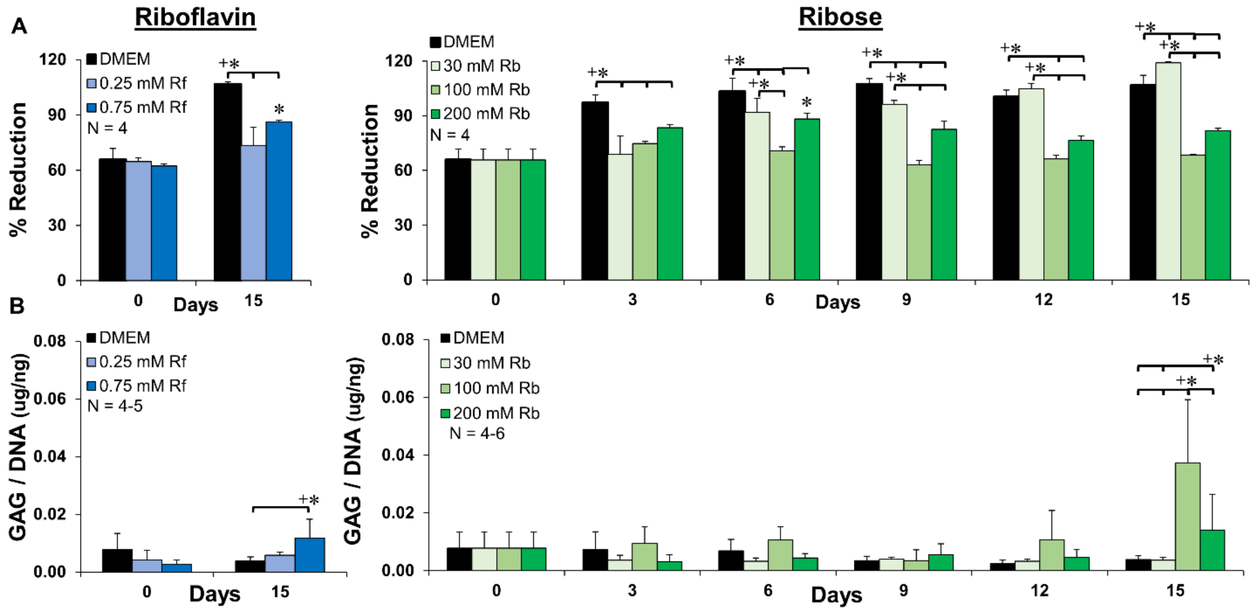


**Supplemental Figure 2:** A) Percent reduction of alamarBlue as a marker of metabolic activity demonstrates reduced reduction of alamarBlue with riboflavin-465 nm treatment by day 15 and reduced reduction of alamarBlue with high concentrations of ribose (100 mM and 200 mM) starting at day 3. B) GAG normalized to DNA demonstrates that both concentrations of riboflavin had little differences in GAG accumulation compared to control constructs throughout culture, while high concentrations of ribose (100 mM and 200 mM) resulted in significantly higher GAG accumulation by day 15. Significance compared to *day 0 of treatment group or + bracketed group ($p$<0.05).

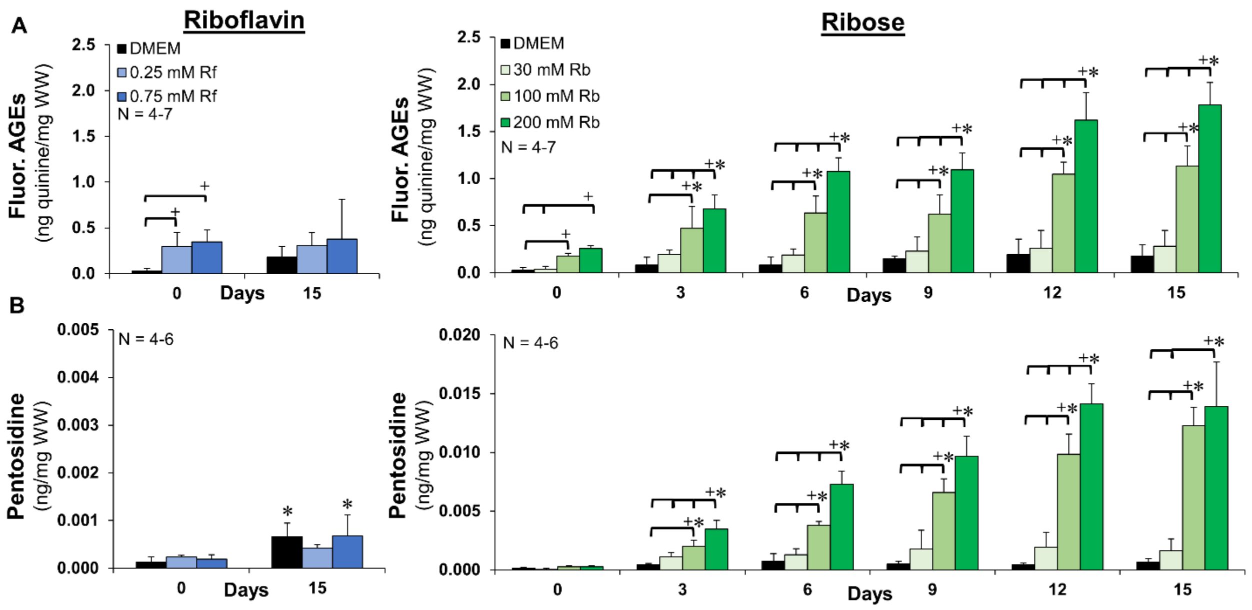


**Supplemental Figure 3:** A) Fluorescent AGEs normalized to wet weight (WW). B) Pentosidine from auto-fluorescence normalized to wet weight (WW). Both concentrations of riboflavin induced a significant increase in total AGEs at day 0, which were maintained through day 15, but had little effect on pentosidine crosslinks. Ribose induced significant increases in total AGEs and pentosidine crosslinks in a dose dependent fashion with time in culture. Significance compared to *day 0 of treatment group or + bracketed group (*p*<0.05).

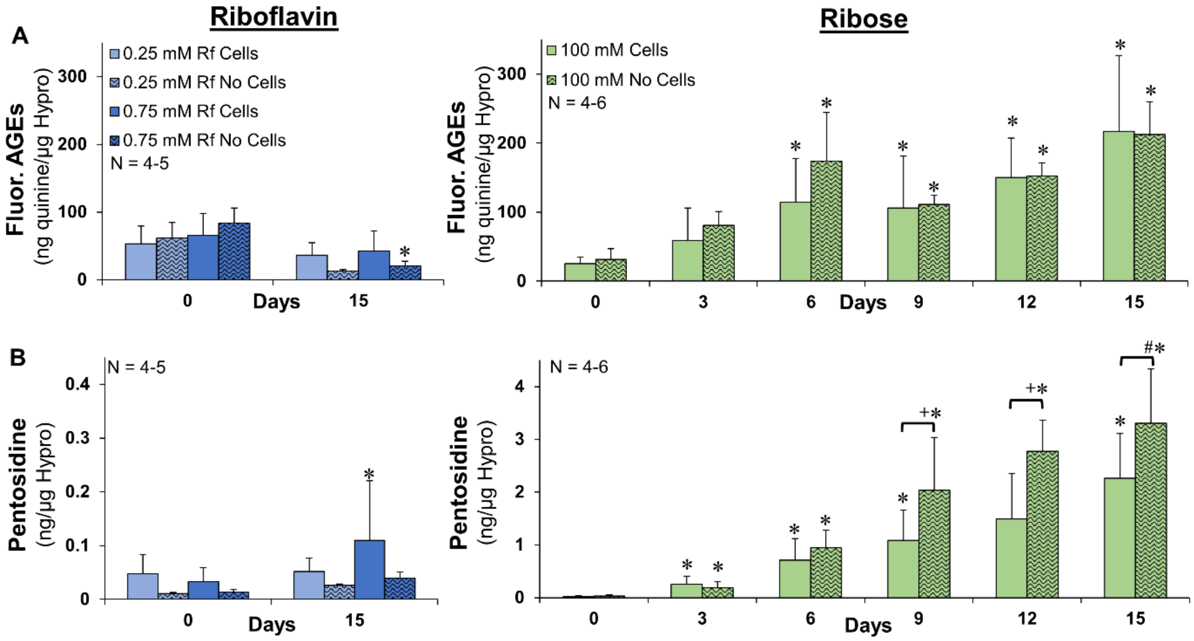


**Supplemental Figure 4:** Induced glycation with and without cells. A) Fluorescent AGEs normalized to collagen content represented by hydroxyproline (hypro) and B) pentosidine from auto-fluorescence normalized to hypro. Both riboflavin-465 nm and ribose treatment induced similar levels of fluorescent AGEs whether collagen gels had cells or did not have cells. The presences of cells had no significant effect on pentosidine accumulation in riboflavin-465 nm induced gels, however cell-seeded constructs did have reduced pentosidine accumulation starting at day 9 in ribose treated constructs compared to acellular gels. Significance compared to *day 0 of treatment group or +bracketed group ($p$<0.05). #trending difference for bracketed group ($p$<0.1)
